# Scaffold-enabled high-resolution cryo-EM structure determination of RNA

**DOI:** 10.1101/2024.06.10.598011

**Authors:** Daniel B. Haack, Boris Rudolfs, Shouhong Jin, Kevin M. Weeks, Navtej Toor

## Abstract

Cryo-EM structure determination of protein-free RNAs has remained difficult with most attempts yielding low to moderate resolution and lacking nucleotide-level detail. These difficulties are compounded for small RNAs as cryo-EM is inherently more difficult for lower molecular weight macromolecules. Here we present a strategy for fusing small RNAs to a group II intron that yields high resolution structures of the appended RNA, which we demonstrate with the 86-nucleotide thiamine pyrophosphate (TPP) riboswitch, and visualizing the riboswitch ligand binding pocket at 2.5 Å resolution. We also determined the structure of the ligand-free apo state and observe that the aptamer domain of the riboswitch undergoes a large-scale conformational change upon ligand binding, illustrating how small molecule binding to an RNA can induce large effects on gene expression. This study both sets a new standard for cryo-EM riboswitch visualization and offers a versatile strategy applicable to a broad range of small to moderate-sized RNAs, which were previously intractable for high-resolution cryo-EM studies.

## INTRODUCTION

There is great interest in the development of therapeutics targeting RNA structures *in vivo* for the treatment of disease. Risdiplam, for which the lead hit was discovered by phenotypic screening, is an example of an FDA approved drug that targets RNA for the treatment of spinal muscular atrophy (SMA) (*1*). The design of small molecule drugs to target proteins is notably accelerated by studying structure activity relationships (SAR) to optimize binding to protein targets, as informed by routine visualization of protein-ligand complexes. However, structure-informed SAR is currently difficult and rare for the development of small molecules targeting RNA structures. Most RNAs do not crystallize readily for x-ray crystallography and also present difficulties for cryo-EM structure determination. Structure determination of protein-free RNAs has proven especially difficult. Routinely obtaining high-resolution structures of RNA and RNA-ligand complexes is crucial for establishing molecular mechanisms of action in various biological contexts. Cryo-electron microscopy (cryo-EM) has revolutionized structural biology with thousands of protein structures solved and shared in the Protein Data Bank (PDB). However, there has only been a single protein-free RNA (Tetrahymena group I intron) determined at 3 Å resolution or better using cryo-EM (*2*). The majority of protein-free RNAs exhibit resolutions ranging from 5 to 10 Å (*3*), suggesting that RNA is not as amenable to cryo-EM structure determination as are proteins. Technologies that would allow for the routine application of cryo-EM to protein-free RNA would have a significant positive impact for RNA structural biology and for the development of small molecule drugs targeting RNA.

RNA presents several difficulties in cryo-EM structure determination. We have found that vitrification on cryo-EM grids often results in the denaturation and aggregation of RNA. These interfering processes result in only a small number of native particles to be visualized, which are insufficient to obtain a high-resolution reconstruction. In addition, the few particles that are visualized often exhibit a preferred orientation that further complicates structure determination because the views of the particle are not sufficiently populated for high resolution 3D reconstruction. Therefore, there is a compelling need for tools and technology to overcome these obstacles and enable routine high-resolution cryo-EM analysis of protein-free RNAs.

X-ray crystallography can also be applied for the structure determination of RNA, but has limitations. It is very challenging to form crystals with protein-free RNAs due to the relatively homogeneous negatively charged surface making it difficult to form unique crystal contacts. It is also likely that RNA conformational dynamics are not fully assessed in a crystal lattice because the constraining crystal lattice packs macromolecules together at concentrations approaching hundreds of mg/ml. In this environment, helical dynamics would be suppressed, and the overall RNA structure is favored to be in a compact state. Our structural studies of group II introns supports this hypothesis. We attempted to characterize branch-site helix dynamics using crystallography and observed only small scale movements in the domain VI helix of a group IIB intron (*4*). We subsequently determined the cryo-EM structure of a homologous group IIB intron, with a shared secondary structure, and found that the branch-site helix engages in a 90° swinging action between the two steps of splicing (*5*). We hypothesize that similar conformational dynamics could be observed for riboswitches if cryo-EM were feasible for these small RNAs.

Riboswitches are RNA structures found in mRNAs that bind small molecule ligands to affect gene expression by altering either transcription or translation of a downstream gene, by either sequestering or exposing a regulatory RNA structure (*6*). Most riboswitches have two domains, an aptamer domain which binds the regulating ligand and forms an RNA motifs with a specific higher-order tertiary structure, and an expression domain whose formation, or not, regulates either transcription or translation of the downstream gene. The riboswitch mechanism implies there is an equilibrium between different conformations, with the presence or absence of ligand shifting this equilibrium to govern the resulting effects upon gene expression (*7, 8*). However, crystal structures of most riboswitches only show minor differences between the bound and apo states (*9–14*). We hypothesize that this is likely due to the constraining environment of the crystal lattice. Obtaining high-resolution cryo-EM structures would be preferable since RNA is frozen in solution, which would allow one to trap alternative conformations.

One approach for solving RNA structures via cryo-EM is to use a scaffold to which one could attach an RNA motif of interest. The rationale for attaching a target RNA to a large scaffold is that the favorable biophysical properties that result in high resolution for the scaffold would propagate to the target RNA. The larger mass of the overall RNA also facilitates particle picking and alignment for 3D reconstruction. This approach has been attempted with the Tetrahymena group I intron, in multiple adaptations (*15, 16*). This group I intron, in isolation, has been solved to ∼3 Å resolution (*2*). However, to date, this approach has resulted in only ∼5 Å resolution for the target RNA (*15, 16*), which does not allow for the discrimination of individual nucleotides and makes precise modeling impossible.

Here, we report our development of a group II intron scaffold that allows for cryo-EM structure determination of attached small RNAs at high resolution. Using this approach, we solved cryo-EM structures of the thiamine pyrophosphate (TPP) riboswitch in both ligand-bound and ligand-free (apo) states. We visualized the ligand-binding pocket of the TPP riboswitch at 2.5 angstroms resolution, which enabled precise modeling of the thiamine pyrophosphate ligand. Comparisons of the ligand-bound and apo structures reveal conformational dynamics that inform a mechanism for riboswitch function. Going forward, this technology allows structure determination at sufficiently high resolution to enable structure-based SAR for small molecules, and drugs, that bind to RNA.

## RESULTS

### Search for a high-resolution RNA scaffold

We sought to identify an improved scaffold to hold small target RNAs for high-resolution cryo-EM structure determination. An ideal scaffold should have the following properties: (1) exhibit a minimal grid orientation preference, (2) have high solubility, (3) have an RNA structure resistant to denaturation, (4) have a molecular mass >100 kDa, and (5) contain a bridging region that can accommodate target RNAs. An RNA with these properties should ideally result in cryo-EM density of 3 Å or better resolution.

We screened multiple group II introns for their suitability as scaffolds for cryo-EM. Group II introns are self-splicing catalytic RNAs that have six distinct domains and typically range in size from 400 to 850 nucleotides. We first tested the attachment of a group IIC intron from *Oceanobacillus iheyensis* (*O.i.*) (*17*) (120 kDa) to a larger (scaffolding) group IIB intron ribonucleoprotein complex from *Thermosynecoccus elongatus* (*T.el.*) (*5*) (300 kDa) to form the *T.el.*-*O.i.* fusion construct. When local refinement was performed on the scaffold portion of the density, a 3.5 Å map of the RNP was recovered (**Fig. 1A**). However, when local refinement was shifted to density corresponding to the embedded IIC intron, the resolution of the target RNA was limited to ∼5 Å. This resolution limit is consistent with prior experiences for scaffold-based approaches (*15, 16*). The RNP scaffold does display severe orientation bias, which is still present after employing a tilted data collection strategy (**Fig. 2C**). However, we were encouraged to observe that the overall fold of the embedded RNA was maintained even though the method resulted in only a moderate resolution map (**Fig. 2B**). We then tested the ability of the group IIC intron in isolation to function as a scaffold. The *O.i.* intron (**Fig. 2A**) folds into a stable structure that crystallizes in a wide variety of precipitants (*17*). However, this particular intron has the unwanted tendency to undergo intramolecular hydrolytic cleavage at multiple sites (*18*), which may also cleave an attached RNA target. Therefore, we utilized an inactive mutant in which an active site residue is mutated to abolish catalytic activity (*19*). In our first attempt, we obtained a reconstruction at 2.4 Å resolution for this group II intron (**Fig. 2B, 2C, and Table S1**). In addition, this intron exhibited a very favorable lack of orientation preference (**Fig. 2D**) and has a stable solvent-exposed stem that could be used to embed an RNA of interest (domain III). The orientation distribution map is evenly populated, which is highly unusual for a protein-free RNA in our experience. Based on these favorable properties, we selected this group IIC intron as a scaffold candidate for the attachment of target RNAs for high resolution cryo-EM.

**Figure 1.**
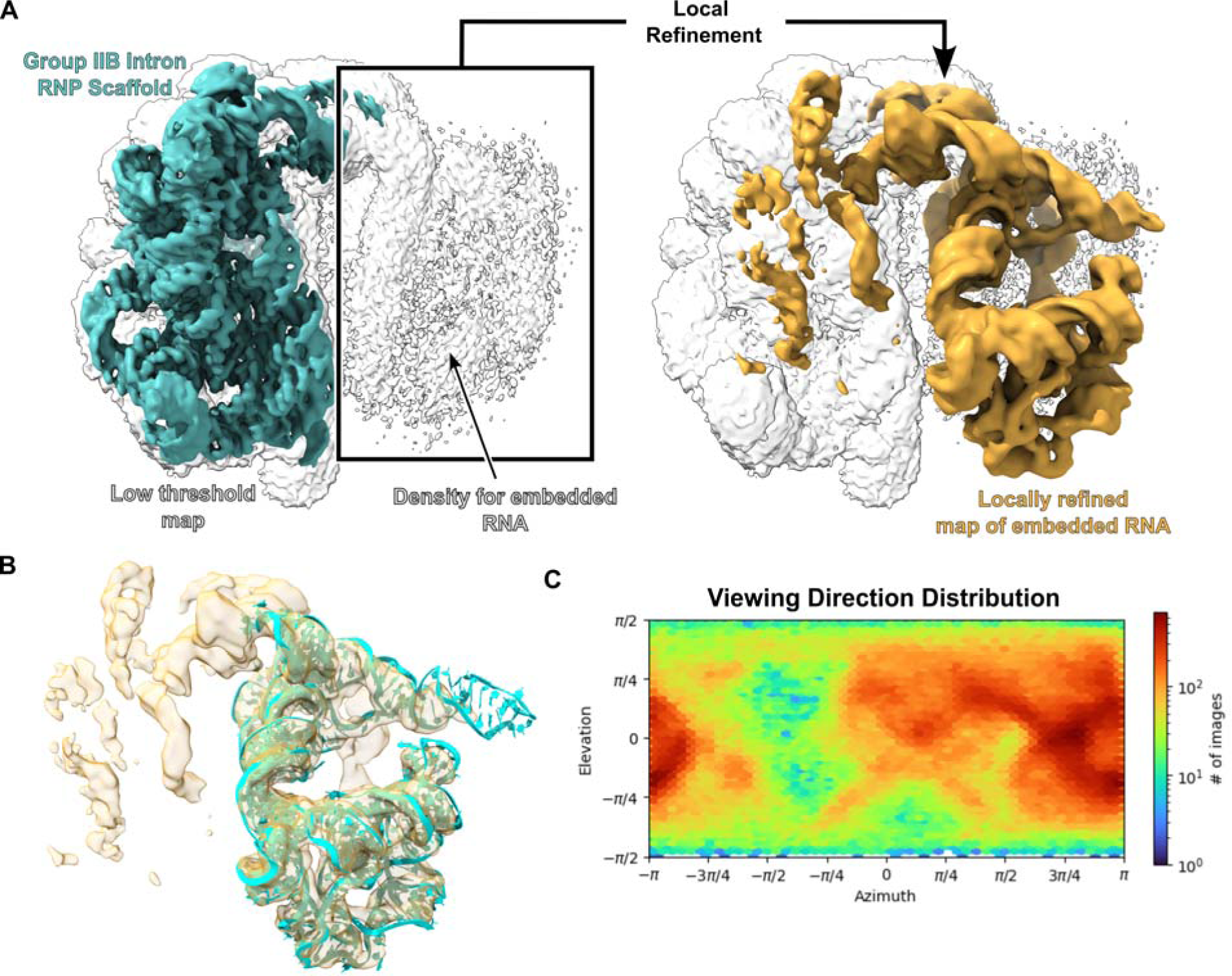
Initial scaffold development using a group IIB intron RNP. **(A)** The maps of the group IIB RNP were generated using a tilted data collection strategy with the stage set to 30°. The map on the left was globally refined and shown at both a high (teal) and low threshold (white). The group II intron RNP displays clear RNA features at high threshold and density for the embedded RNA is observed at low threshold. Performing local refinement on the density corresponding to the embedded RNA yielded the map on the left (gold) which has been overlaid on the low threshold globally refined map (white) for clarity. **(B)** A model for the embedded RNA was fit into the locally refined map. The fit shows that the overall fold of the embedded RNA was maintained. **(C)** The viewing direction distribution for the data set is shown. Even with a 30° stage tilt, there is still a preferred orientation for this specimen.

**Figure 2.**
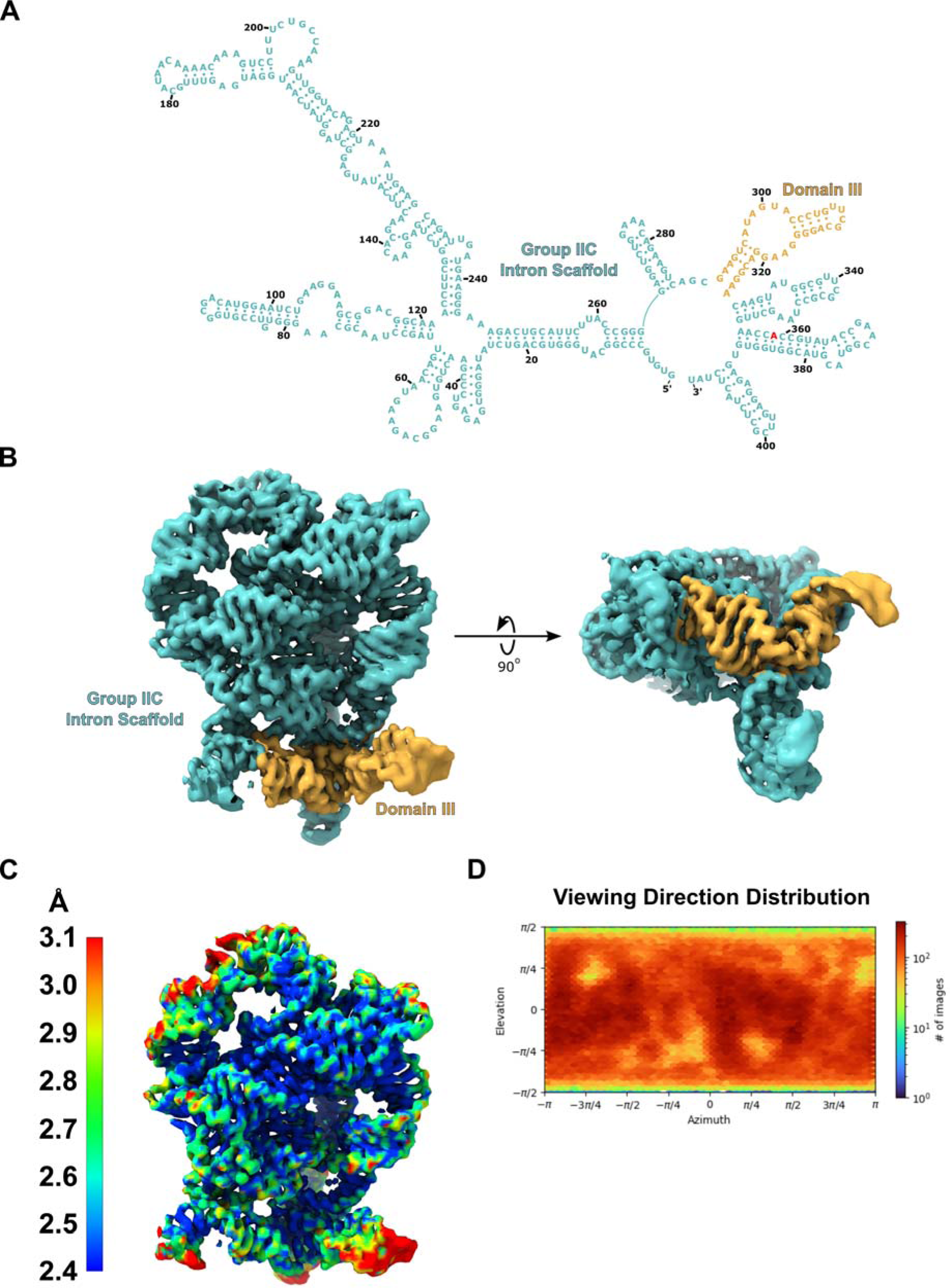
Identification of a high-resolution group IIC intron scaffold. **(A)** The secondary structure of the *O.i.* group IIC intron scaffold (teal) is shown highlighting domain III (gold). The mutation to the G to A mutation to the active site is highlighted red. **(B)** Cryo-EM density is shown for the scaffold. Domain III (gold) forms a stable stem loop that is extruded away from the rest of the folded RNA. **(C)** A local resolution map shows that the core of the scaffold has a resolution of 2.4 Å. **(D)** The viewing direction distribution plot generated from cryoSPARC is shown. The plot is evenly populated indicating that the group IIC intron scaffold exhibits no preferred orientation.

### TPP riboswitch as an RNA target

We attached the aptamer domain of the thiamine pyrophosphate (TPP) riboswitch (*20*) (86-nt) to our group II intron scaffold (*O.i.*-TPP). The TPP riboswitch binds to thiamine pyrophosphate and modulates gene expression, with a wide phylogenetic distribution in both prokaryotes and eukaryotes. Multiple crystal structures of this riboswitch in the presence and absence of thiamine pyrophosphate revealed little difference in the overall RNA fold (*21–23*). A crystal structure of a TPP riboswitch crystallographic dimer was recently solved in the absence of the ligand (*24*). However, in this study, the density did not match the small-angle x-ray scattering reconstruction performed in solution, again emphasizing that crystal packing interferes with the sampling of conformations found natively in solution.

### High resolution reconstruction of the TPP riboswitch

High-resolution cryo-EM typically could not be applied to the TPP riboswitch, given its small size of 86 nucleotides (27.5 kDa). The TPP riboswitch was covalently attached to the solvent exposed domain III stem of the group IIC intron through a rigid helix (**Fig. 3A**). The domain III extension is not phylogenetically conserved (*25*) and can tolerate sequence insertions without affecting the overall intron fold. The resulting fusion construct is 493 nucleotides in length and was synthesized using *in vitro* transcription with subsequent native purification to avoid RNA misfolding. SHAPE-MaP (*26*) confirmed that the secondary structure of the linked group II intron and riboswitch structure maintained the expected fold for both RNA domains (**Fig. 3A and S1**). SHAPE-MaP data also confirmed binding of the TPP ligand to the embedded riboswitch and revealed expected local conformational changes in the riboswitch (*27*). These ligand-induced changes included modest rearrangement of the P2 and P3 helices, and reduced reactivities in the P3-L5 region and in the J3-2 and J2-4 elements. Cryo-EM data collection and subsequent processing were performed for the scaffold attached to the riboswitch (**Fig. S2**). Clear additional signal, corresponding to the attached riboswitch, was visible in all 2D class averages as compared to the 2D classes of the scaffold alone (**Fig. 3B**). Local refinement focused on the scaffold yielded a 3D reconstruction with a resolution of 2.4 Å for the core of the intron (**Fig. 3C**). A comparison of the resulting maps and models for the scaffold alone and with the TPP riboswitch embedded shows that the attachment process did not affect the overall fold of the scaffold **(Fig. S3)**. We then performed local refinement on the riboswitch component, which yielded a reconstruction with a global resolution of 3.1 Å and 2.5 Å for the ligand-binding pocket (**Fig. 3D)**. The viewing distribution plot shows that embedding the riboswitch did not create a preferred orientation for the overall fusion construct (**Fig. 3E**). This resolution yielded clear density for the functional groups in this small molecule and enabled modeling of the TPP ligand with high confidence. Density for three magnesium ions, coordinated to the pyrophosphate moiety, are clearly visible (**Fig. 4**). In previous crystal structures of this riboswitch (*22, 23*), only two metal ions are visualized binding to the pyrophosphate. These magnesium ions are essential for binding of TPP to this riboswitch. The cryo-EM structure of this ligand-bound riboswitch exhibits the classic closed conformation observed in the crystal structures (**Fig. S4**).

**Figure 3.**
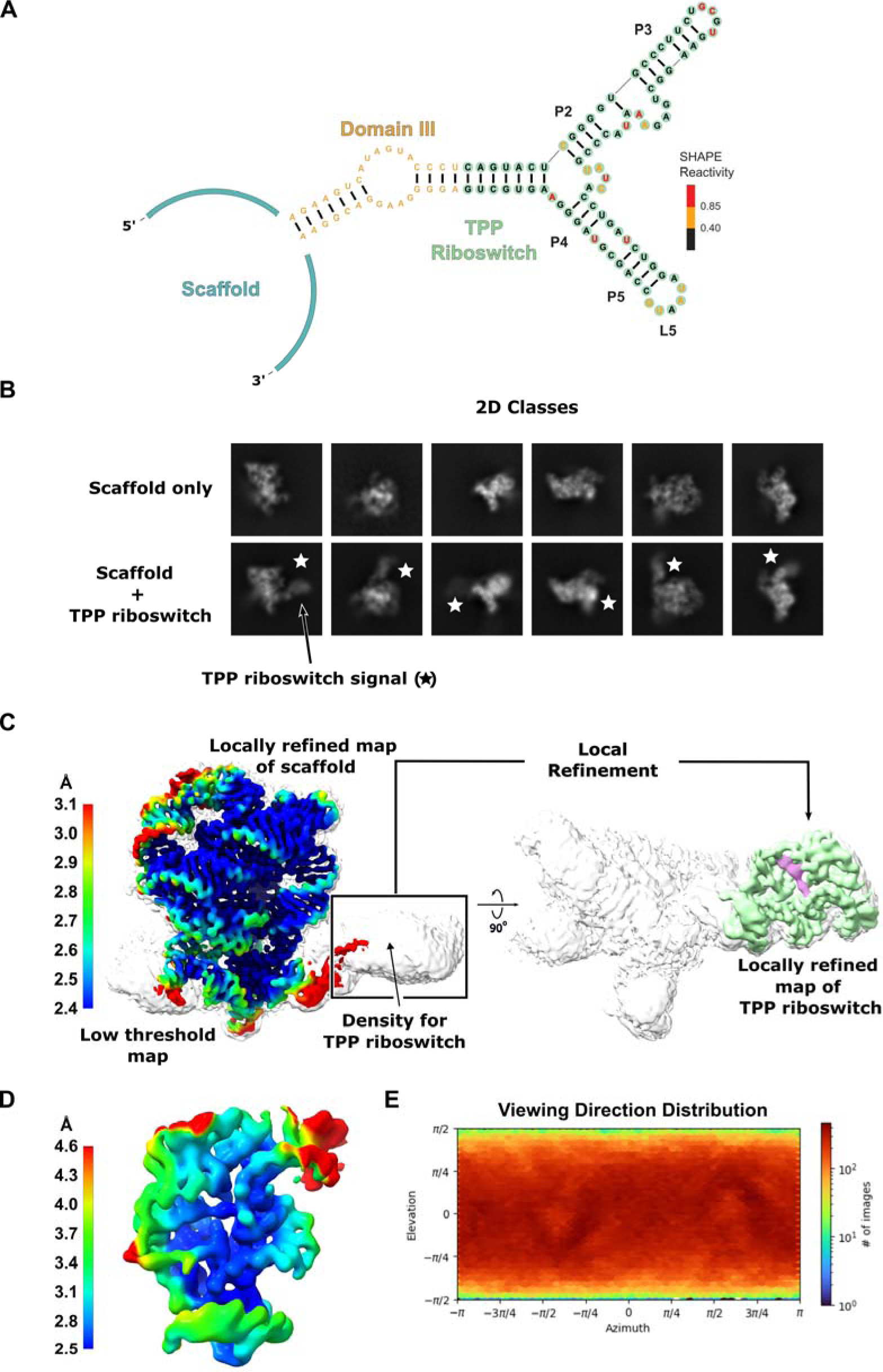
High-resolution cryo-EM structure determination of the TPP riboswitch using the group IIC intron scaffold. **(A)** A secondary structure of the TPP riboswitch (green) attached to the scaffold through the stem of Domain III (gold) of *O.i.* is shown. Nucleotides in the TPP riboswitch aptamer domain are colored by SHAPE reactivity. Experiments were performed in the presence of ligand; for full SHAPE data, see Fig. S1. **(B)** 2D class averages of the scaffold alone and attached to the TPP riboswitch are shown. Signal corresponding to the TPP riboswitch (star) is clearly visible in the 2D classes. **(C)** A local resolution map of the scaffold (left) on was generated from focused refinement and overlaid on a low threshold globally refined map (white). The locally refined map of scaffold maintains high-resolution features after the TPP riboswitch is embedded. Density for the TPP riboswitch is visible in the globally refined map at low threshold. A locally refined map of the TPP riboswitch (green) is shown on the right and overlaid on the same low threshold globally refined map (white) for clarity. **(D)** A local resolution map for the TPP riboswitch. The resolution surrounding the TPP binding site is 2.5 Å. **(E)** A viewing direction distribution plot is shown for the globally refined scaffold TPP data. The scaffold does not show any orientation preference with the added sequence of the riboswitch.

**Figure 4.**
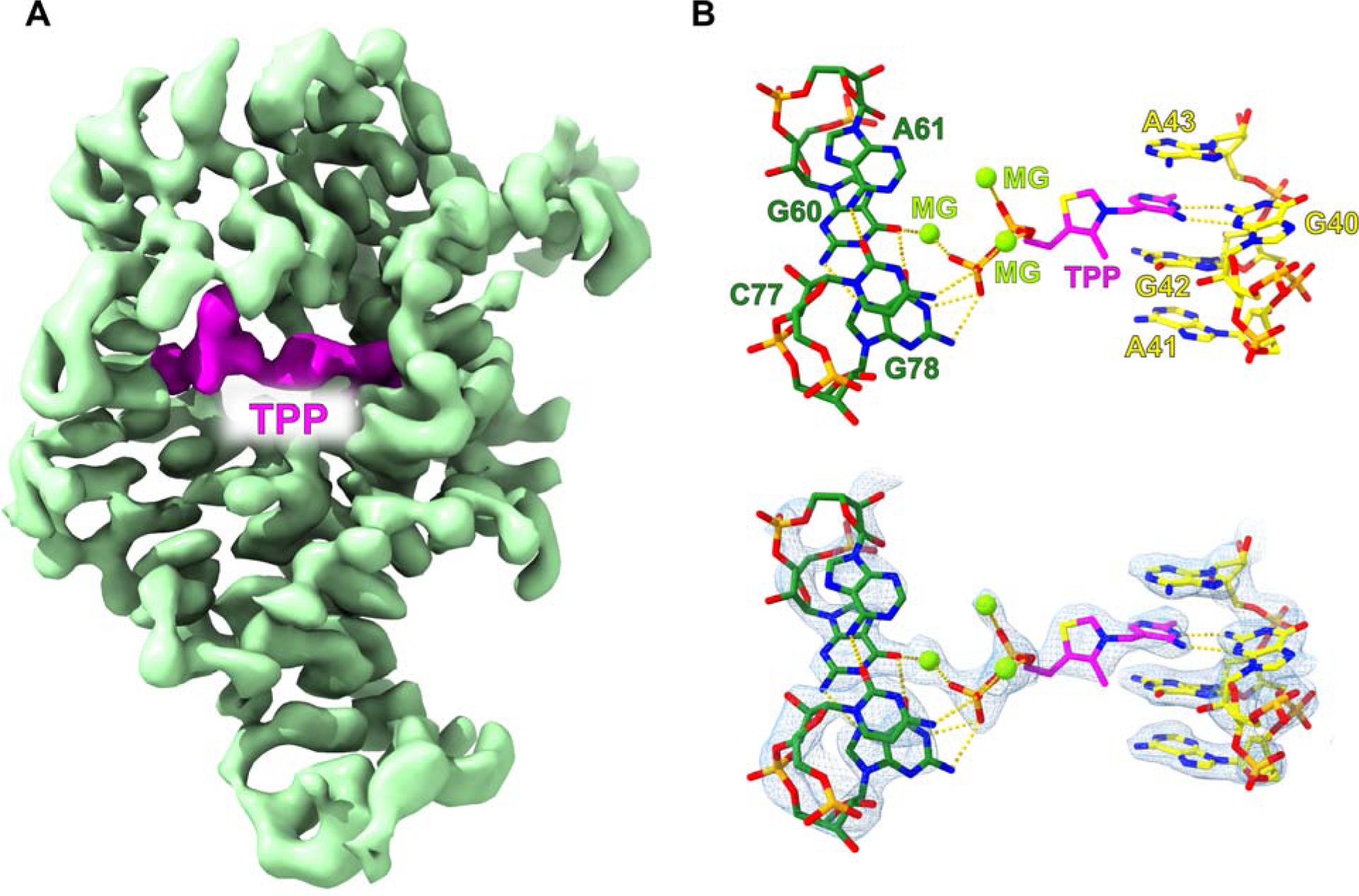
High-resolution cryo-EM structure of the TPP riboswitch. **(A)** A sharpened map of the TPP riboswitch (green) is shown on the left with clear density for a bound thiamine pyrophosphate ligand (magenta). **(B)** A model was built and refined into the cryo-EM density. The high-resolution density surrounding the thiamine pyrophosphate binding site allowed for the precise characterization of ligand binding by the riboswitch. The model overlaid with density is shown below.

In contrast, focused refinement of the ligand-free riboswitch revealed an open “Y” conformation at ∼6 Å resolution (**Fig. 5A**). The TPP aptamer structure can be subdivided into two domains, comprised of thiamine-sensing and pyrophosphate-sensing helices. In the ligand-free (apo) structure, the thiamine-sensing and pyrophosphate-sensing domains form a ∼90° angle. In contrast, upon ligand binding, the RNA adopts a compact tertiary structure in which the two stems are parallel to each other and the P3 and L5 motifs form a long-range tertiary interaction, stabilizing the binding pocket for TPP. The apo state is also more dynamic due to the absence of the ligand and absence of stabilizing tertiary interactions. The thiamine sensing-stem adopts an alternate stable conformation in the apo state. Major differences are two-fold: the J2-4 element and the pyrophosphate-sensing stem are highly disordered, and the thiamine-sensing stem adopts a distinct conformation in which there is clear density for an additional helical groove that extends this helix compared to the bound state (**Fig. 5A**). These substantial conformational differences, visualized directly here, support a molecular model that rationalizes changes in the riboswitch structure as visualized by in-solution SHAPE probing of the ligand-free state (Fig. S1 and ref. 28) (*28*).

**Figure 5.**
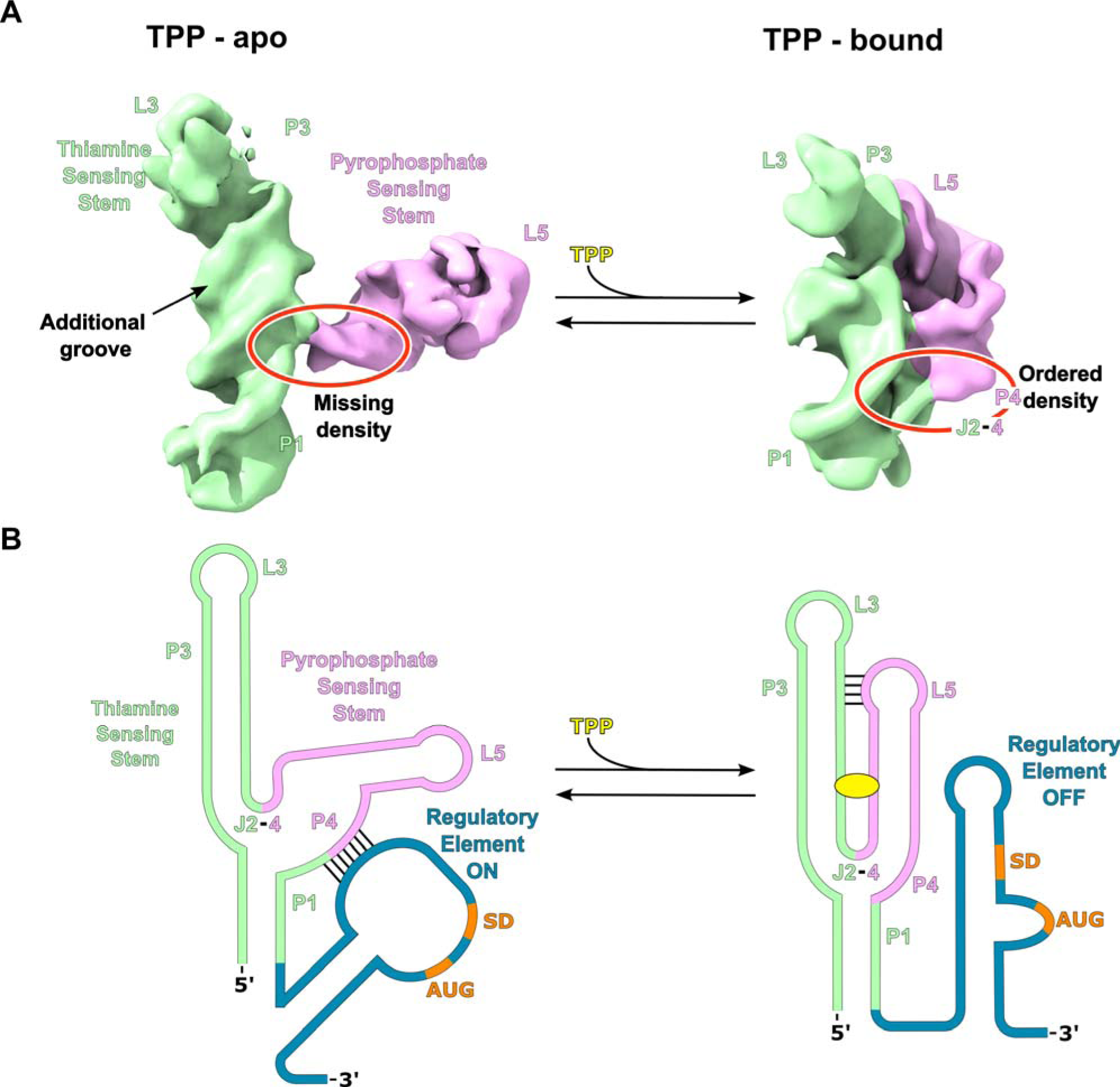
A model for translational regulation by the TPP riboswitch. **(A)** Density for the apo and TPP bound states of the TPP riboswitch are shown. The TPP bound map was low-pass filtered to 6 Å. The P3-L5 interaction is disengaged in the apo state (left) allowing the pyrophosphate sensing stem (purple) to rotate causing the riboswitch to adopt an overall Y shaped conformation. In the apo state the base of the L5 stem loop is largely disordered with portions of P1, J2-4, and P4 missing density completely (circled red). In contrast, the thiamine sensing stem (green) appears to have an alternate stable conformation in the apo state with the appearance of an additional minor groove. When TPP is bound (right), the P3-L5 interaction is engaged, and the riboswitch compacts leading to ordered P1, J2-4, and P4 regions (circled red) properly forming the three-way junction. **(B)** The above structural information provides insights into the mechanism of translational regulation by the TPP riboswitch. In the apo state, the base of the L5 stem loop is disordered allowing it to interact with the downstream regulatory element (blue) through complementary base pairing. This interaction causes a rearrangement of the regulatory element structure making the Shine-Dalgarno sequence (SD, orange) accessible for the ribosome to initiate translation. The thiamine sensing stem having a stable conformation in the apo state likely promotes the formation of this L5 stem loop/regulatory element interaction. In the bound state, the equilibrium is shifted to the compacted state with an ordered L5 stem loop which prohibits the interaction of the L5 stem loop with the regulatory element. This ultimately leads to a loss in ribosome initiation and translation.

The dynamic nature of the unbound riboswitch explains why this state has not previously been captured using x-ray crystallography. The resolution of the apo state is limited due to the inherent dynamics of this destabilized RNA, yielding fewer particles going into the 3D reconstruction. Critically, these two structures represent a basis for modeling the transition through which the riboswitch modulates gene expression upon ligand binding. In this model, the opening of the riboswitch allows it to interact with the downstream regulatory element and affect either translation or transcription through an alteration in RNA structure. Our study provides the first cryo-EM data showing large-scale conformational changes in a riboswitch upon ligand binding, and which supports a specific mechanistic model for changes in gene expression.

Several properties conferred by the scaffold appear to contribute to the high quality of the target RNA reconstruction. The scaffold-riboswitch construct shows a uniform orientation distribution map (**Fig. 3E**), similar to that observed for the scaffold alone. The orientation properties of the scaffold alone in vitrified ice are carried over to the fusion construct with the riboswitch. In addition, the scaffold folds stably which likely protected the riboswitch from denaturation at the air-water interface. The scaffold produced easily visible, high contrast particles within the vitrified ice, facilitating particle identification and improving the accuracy of initial particle alignment. The riboswitch is rigidly attached to the scaffold via stable linking helix, allowing accurate particle alignments to be propagated to the embedded RNA.

## DISCUSSION

We have developed a technology for high-resolution structure determination of small RNAs by fusing these RNAs with a group II intron scaffold. This scaffold strategy is the first to allow cryo-EM structure determination of a target RNA to better than 3 Å resolution. In the case of the TPP riboswitch, the resolution observed here is the highest ever achieved via cryo-EM for an RNA less than 100 nucleotides. The cryo-EM structure of the TPP riboswitch reveals the ligand-binding pocket at 2.5 Å resolution, sufficient to enable de novo modeling of the thiamine pyrophosphate ligand (MW 425 g/mol). The predominant ligand-free state of the riboswitch reveals a large-scale conformational change in which that the two helices forming the binding pocket move apart to form an overall Y-shaped fold. There is a significant difference in the overall fold of the thiamine sensing stem between the ligand-free and bound states. The thiamine sensing stem in the ligand-free state has a more elongated helix with the appearance of an additional helical groove (Fig. 5A). This is consistent with the rearrangement of the secondary structure in the base of this stem to form additional pairing interactions. It is likely that a rearrangement in secondary structure is required for the interaction with the pyrophosphate group upon ligand binding.

The large conformational difference between the bound and apo forms supports a mechanism for riboswitch function. The native TPP riboswitch features a downstream regulatory element consisting of a stem loop containing the start codon of the mRNA. In this model, the ligand-free riboswitch initially exists in a relaxed, dynamic Y-shaped form and interacts with the regulatory stem loop to render the start codon accessible for translation initiation (**Fig. 5B**). Once the riboswitch binds the ligand, the riboswitch clamps down and forms a compact structure that disengages the riboswitch from the regulatory stem loop, creating new structures that sequester the start codon and inhibit translation. Large conformational changes, of similar magnitude upon ligand binding, are likely a general mechanism for riboswitch function.

This work shows that it is possible to obtain high resolution structures of small RNAs using cryo-EM. We anticipate that this scaffolding approach will allow for routine structure determination of many biologically significant RNAs. The ability to visualize a small molecule bound to RNAs with complex structures (*29*) using cryo-EM now creates a broad opportunity to use this approach in drug discovery efforts targeting RNA structures.

## MATERIALS AND METHODS

### Plasmid cloning

The mutant *O.i, O.i.-*TPP, and *T.el*4h-*O.i.* genes were synthesized (Genscript) and cloned into a pUC57 vector using the EcoRV restriction site. The cloned plasmids were transformed into DH5α cells. For all cryo-EM experiments, the genes contained a 4-nt 5′ exon and a 3-nt 3′ exon followed by a BamHI cut site. The *T.el.*4h maturase expression plasmid used in this study was previously described (*5*).

### *In vitro* RNA transcription and purification

The *O.i, O.i*-TPP, and *T.el*4h-*O.i.I* plasmid was linearized using an engineered (BamHI for *O.i.* and *O.i.*-TPP and HindIII for *T.el*4h-*O.i.*) restriction site (NEB). 50 μg of template DNA was added to a total volume of 1 mL of *in vitro* transcription buffer (50 mM Tris-HCl pH 7.5, 25 mM MgCl_2_, 5 mM DTT, 2 mM spermidine, 0.05% Triton X-100, and 5 mM of each NTP). T7 polymerase and thermophilic inorganic phosphatase was added to begin RNA synthesis. The reaction mixture was incubated at 37°C for 3 hrs. CaCl_2_ was added to a final concentration of 1.2 mM along with Turbo DNase and placed at 37°C for 1 hr to fully digest the DNA template. Proteinase K was subsequently added and incubated at 37°C for an additional hour. The resulting solution was centrifuged to remove any precipitate and then filtered through a 0.2 μm filter. The filtered solution was buffer exchanged a total of 7 times, each time using 14 mL of filtration buffer (5 mM Na-cacodylate pH 6.5 and 4 mM MgCl_2_) and a 100 kDa molecular weight cut-off filter. After the final buffer exchange step, the RNA was concentrated to approximately 10 mg/mL for use in downstream cryo-EM experiments.

### SHAPE-MaP RNA structure probing

SHAPE-MaP assays were performed essentially as described previously (*30*). Briefly, 10 pmol RNA in 19 µL probing buffer (300 mM HEPES, pH 8.0, 2 mM MgCl2, 40 mM NaCl) was mixed with 1 uL of TPP (final concentration, 50 µM). After incubation for 15 min at 37 °C, 9 uL of RNA-ligand solution was added to 1 uL 1M 2A3 (*31*) and mixed thoroughly. After incubation for 20 min, the reaction was quenched by addition of 5 uL of 1M DTT. No ligand reactions were conducted in parallel. RNA was purified (G-25 spin column) and subjected to reverse transcription (SuperScript II, Invitrogen) using a heat-cool protocol [25°C for 10 min, 42°C for 90 min, 10 cycles of (50°C for 2 min, 42°C for 2 min), 70°C for 10 min]. DNA libraries were prepared from cDNA using a two-step PCR process (Q5 Hot Start High-Fidelity DNA Polymerase, New England Biolabs). PCR 1 used Step-1 Forward primer and Step-1 Reverse primer, and performed for 20 cycles; PCR 2 used Universal Forward primer and Universal Reverse prime, and performed for 10 cycles. All cDNA or dsDNA were purified (Mag-Bind TotalPure NGS beads, Omega), quantified (Qubit dsDNS dsDNA Quantification Assay Kits, Life Technology), and visualized to confirm integrity (2100 Bioanalyzer Instrument, Agilent). The libraries were sequenced on a MiSeq system (Illumina). Mutations were aligned and parsed using ShapeMapper (v2.15) (*32*) using default parameters.

RT primer sequence: 5’-CTAATAGAGT AGAGCGAACT CCTCTC-3’.

Step-1 Forward primer: 5’-CCCTACACGA CGCTCTTCCG ATCTNNNNNT TATGTGTGCC CGGCATG-3’. Step-1 Reverse primer: 5’-GACTGGAGTT CAGACGTGTG CTCTTCCGAT CTNNNNNCTA ATAGAGTAGA GCGAACTCCT-3’.

Universal Forward primer: 5’-AATGATACGG CGACCACCGA GATCTACACT CTTTCCCTAC ACGACGCTCT TCCG-3’. Universal Reverse primer: 5’-CAAGCAGAAG ACGGCATACG AGAT (8nt-barcode) GTGACTGGAG TTCAGAC-3’.

### Group IIB intron RNP assembly

The RNP used in this study was assembled and purified as previously described (*5*).

### EM sample preparation

For all grids prepared with TPP bound, 100 uL of 2 mg/mL RNA was incubated with 1 mM thiamine pyrophosphate at room temperature for 15 minutes prior to freezing. For all grids prepared with the riboswitch in the apo state, this step binding step was omitted. To freeze the grids, 3.5 μL of freshly prepared RNA sample at 2.0 mg/mL was applied to a glow discharged (40 mBar, 15 mA for 30 s using a PELCO easiGlow) copper R1.2/1.3 300-mesh grid (Quantifoil). The grid was blotted with a filter paper (Whatman No.1) at 4 °C in a cold room before plunging frozen into liquid ethane/propane (37.5/62.5) mix using a manual plunger. The *T.el*4h-*O.i* RNP specimen, 3.5 μL of freshly prepared RNP at 1 mg/mL was applied to a glow discharged (40 mBar, 15 mA for 120 s using a PELCO easiGlow) UltrAuFoil R1.2/1.3 300-mesh grid (Quantifoil). The grid was blotted with a filter paper (Whatman No.1) at 4 °C in a cold room before plunging frozen into liquid ethane/propane (37.5/62.5) mix using a manual plunger.

### Cryo-EM data collection and processing

Movies were collected on the same Titan Krios microscope (Thermo Fisher) operating at 300 keV, equipped with a K3 Gatan direct electron detector, and were collected with a magnification of 105,000x for a physical pixel size of 0.811 Angstroms and camera operating in super-resolution mode (0.4055 Angstroms/pixel). Movies collected for *O.i.*, *O.i.*-TPP, and *O.i.*-TPP-Apo were collected as 50 dose-fractionated frames with a dose rate of 8 e-/pixel/s for a total exposure time of 4.5 s and total dose of ∼50 e-/Å2. The *O.i.* and *O.i.*-TPP datasets were collected with a defocus range of −0.5 μm to −1.5 μm, whereas the *O.i.*-TPP-Apo dataset was collected with a defocus range of −0.5 μm to −0.9 μm. The *T.el.*-*O.i.* dataset was collected with the same parameters described previously (Ref) as reported in table S1, with the exception of a defocus range of −0.8 μm to −1.9 μm, and an exposure time of 5.4 s. All Movies were semi-automatically collected using SerialEM. 5724 (*O.i.*), 30,314 (*O.i.*-TPP), 15,876 (*O.i.*-TPP-Apo) and 2596 (*T.el.*-*O.i.*) movies were collected with these parameters.

For the *O.i.* dataset, 5,724 movies were collected and imported into Relion 3.1 (*33–35*). Super resolution movies were binned by two and motion-corrected using MotionCor2 (*36*). These motion-corrected micrographs were imported into CryoSPARC 3.3.2 (*37*), and CTF was estimated using Patch CTF. Micrographs were visually inspected and 343 micrographs were thrown out due to poor particle distribution and ice contamination. Particles were picked on the remaining 5,381 micrographs using a general model from Topaz 0.2.5 (*38*) as implemented in CryoSPARC. These ∼1.6 million particles were extracted with a box size of 288 pixels downsampled to 64 pixels, for a final pixel size of 3.6 A/pixel. Particles were ran through 2D classification, and a small subset of particles was selected for ab initio model building. This model was then used for four subsequent rounds of 2-class heterogeneous refinement to classify the ∼1.6 million particles into a 3D class that looked like the group II intron structure. The resulting 725,736 particles were extracted as 320 pixel boxed particles downsampled to 160 pixels for a pixel size of 1.6 A/pixel. The particles were then subjected to several additional rounds of 3D classification, refinement, and CTF refinement, before the final set of 607,776 particles were extracted with a box size of 360 pixels downsampled to 240 pixels, for a final pixel size of 1.22 A/pixel. These ∼600,000 particles were put through a single round of 3D classification with alignment (562,162 particles selected), 2D classification (458,691 particles selected), and 9-class heterogeneous refinement (382,753 particles selected) for Non-uniform refinement. This map refined to a final GSFSC resolution of 2.62 A as reported in cryoSPARC.

For the *T.el.*-*O.i.* dataset, 2596 movies were collected at a 30-degree stage tilt and imported into cryoSPARC 4.2.1. Super resolution movies were binned by two and motion-corrected and CTF was estimated using Patch Motion Correction and PatchCTF, respectively. 1775 motion corrected micrographs were selected for further processing with good particle distribution and little ice contamination. 100 micrographs were randomly selected for initial particle picking using blob picker, and these ∼60,000 particles were extracted for 2D classification. Good 2D references corresponding to ∼22,000 particles were selected for ab initio model building and refinement. The 3D reference from this dataset was used for picking ∼633,000 particles on the full 1775 micrographs, 502,500 of which were then extracted at a box size of 512 pixels downsampled to 128 pixels, for a final pixel size of 3.2 A/pixel. The 3D volume generated by the initial refinement was used as the starting template for three rounds of consecutive 2-class heterogeneous refinement, and the resulting 331,439 particles were put through refinement and extracted with a box size of 512 pixels (0.811 A/pixel). The 322,683 particles were subjected to duplicate removal (20 Angstrom separation), another round of 2-class heterogeneous refinement, refinement, CTF refinement, and 2D classification, for a final set of ∼283,000 particles. The GSFSC (0.143) global resolution of this particle set after non-uniform refinement was 2.88 A as reported by cryoSPARC. These particles were then subjected to hetero-refinement and 3D classification to select particles where the embedded *O.i.* insert was fully intact. The 34,822 particles selected from classification were used for masked local refinement around the *O.i.* insert, which had a GSFSC resolution of 4.02 A as reported in CryoSPARC.

For the *O.i.*-TPP-Apo dataset, 15876 super resolution movies were collected and imported into cryoSPARC 4.2.1. Movies were binned by two, and motion-corrected and CTF estimated using patch motion correction and PatchCTF, respectively. Exposures with CTF resolution fits above 10 Å, relatively thick ice, and high values for the predicted stage tilt angle, were removed, leaving 11,485 micrographs for further processing. Due to difficulties with initial blob picking, a 1,800 micrograph subset was used for template picking using an *O.i.* map processed from a previous dataset. These ∼800,000 particles were used for two rounds of 3-class hetero-refinement, with the final 93,000 particles used to generate a new template for particle picking against the full ∼11,000 micrograph dataset. The ∼8,000,000 particles picked were extracted with a 336 pixel box downsampled to 84 pixels (3.2 A/pixel). These particles were subjected to four consecutive rounds of hetero-refinement (4 classes, 2-class, 2-class, and 2-class). The resulting 1,212,955 particles were then refined and re-extracted to a box of 128 pixels (2.3 A/pixel). Duplicate particles (50 angstrom radius) were removed, and remaining particles were used for a 5-class hetero-refinement. The 1,056,211 particles selected were refined, and extracted with a 256 pixel box (1.27 A/pixel), and then put through another round of refinement and CTF refinement. These 1,032,440 particles were then ran through a 10-class hetero-refinement, and 2 classes that corresponded to the highest resolution densities were selected. These ∼600,000 particles were then subjected to multiple rounds of refinement and CTF refinement to improve resolution, with one final 2D classification job used to remove particles with poor 2D classes. These 547,101 particles were re-extracted with a box size of 448 pixels downsampled to 288 pixels (1.26 A/pixels) and non-uniformly refined to a final global GSFSC resolution of 2.78A as reported by cryoSPARC.

For the apo-TPP embedded structure, those ∼547,000 particles were put through multiple rounds of 3D classification and hetero-refinement to remove conformations for the P2 and P3 stems that did not have resolvable densities, or corresponded to conformations that were not relevant to the TPP riboswitch. Classes that had intact density for an open conformation were pooled together, for a final particle set of 106,285 particles. In parallel, a mask around the *O.i.* scaffold was generated for local refinement of the scaffold, and these 3D coordinates were used for signal subtraction of the scaffold from the particle images. Then, the 3D coordinates of the ∼106,000 particles after non-uniform refinement were used as a starting point for local refinement of the apo-TPP riboswitch in its open conformation. Gaussian priors of an 8 degree rotation and 4 A shift were used for masked local refinement of the apo state, which had a final GSFSC of 4.84 Angstroms as reported in CryoSPARC.

The basic workflow for the high resolution reconstruction of the TPP-riboswitch is outlined in figure S2. In brief, three separate data collections of the *O.i.*-TPP construct were collected (7763 movies, 6303 movies, 16248 movies) with identical parameters, and pooled together as 30,314 super resolution movies. These movies were initially imported into cryoSPARC 4.2.1 for validation and a first reconstruction, and used a similar workflow as that implemented for the *O.i.*-TPP-apo construct. The first maps from cryoSPARC had a resolution close to 3.7 A/pix, and so the super resolution movies were imported into Relion 4.0 as separate exposure groups, and put through a standard workflow of particle picking, 3D classification, refinement, and particle polishing at the end. These particles were then exported into cryoSPARC, where they underwent the process of resolution improvement through non-uniform refinement and particle defocus refinement for a final GSFSC resolution of 2.53 A. Masked refinement of the scaffold was then used for signal subtraction from the polished particles, and the signal subtracted particles were recentered on the density for the TPP riboswitch. This TPP riboswitch density was locally refined with a Gaussian prior of a 3 degree rotation and 3 Å shift, with a final reported GSFSC resolution of 2.96 Å in cryoSPARC.

### Model building and structure refinement

As a starting point for model building for *O.i*., 4DS6 structure coordinates were refined in real space by PHENIX (*39, 40*). The same process was repeated for the locally refined *O.i.*-TPP focused on the scaffold. For the locally refined map focused on the TPP riboswitch, 2GDI structure coordinates were real space refined by PHENIX. Any nucleotide that need to be remodeled was done in COOT (*41, 42*) using the RCrane plugin (*43, 44*). UCSF Chimera was used to make figures depicting the cryo-EM density (*45*). All software was compiled by SBGrid (*46*).

### Quantification and Statistical Analysis

RNA concentrations were determined using a Nanodrop spectrophotometer (Thermo-Fisher). Maturase protein concentrations were determined using an SDS-PAGE gel with a titration of BSA (Thermo-Fisher). To calculate per-reside Full RMSD values, the two models being compared were superposed in COOT using LSQ using the entire sequence for alignment. The superposed models were opened in UCSF Chimera for evaluation using Match -> Align. All map and model validation and statistics were done in PHENIX (Table S1).

## Acknowledgements

We would like to thank the Cal-Cryo Facility at UC Berkeley and Dan Toso for assistance with data collection.

## Author contributions

Conceptualization: DBH, BR, NT

Methodology: DBH, BR, SJ, KMW, NT

Investigation: DBH, BR, SJ, KMW, NT

Visualization: DBH, BR, SJ, KMW, NT

Supervision: KMW, NT

Writing—original draft: DBH, BR, NT

Writing—review & editing: DBH, BR, KMW, NT

## Data and materials availability

All data needed to evaluate the conclusions are available in the main text or the supplementary materials. Structure coordinates have been deposited in the Protein Data Bank (PDB) under accession numbers 9C6I (*O.i.*), 9C6J (*O.i.*-TPP focused on *O.i.*), 9C6K (*O.i.*-TPP focused on TPP). Cryo-EM density maps have been deposited in the Electron Microscopy Data Bank (EMDB) under accession numbers 45247 (*O.i.*), 45248 (*O.i.*-TPP focused on *O.i.*), 45249 (*O.i.*-TPP focused on TPP), 45250 (*O.i.*-TPP apo focused on TPP), and 45251 (*T.el.*-*O.i.* focused on *O.i.*).

