## Supplementary Information for "Scaffold-enabled high-resolution cryo-EM structure determination of RNA"

**Supplementary Figure Legends**

**Figure S1. SHAPE-MaP probing of scaffold-TPP riboswitch construct in the absence and**

**presence of TPP ligand.** (A) SHAPE reactivity profiles in the absence or presence of TPP ligand. TPP riboswitch region is emphasized with green box. Riboswitch structural landmarks are highlighted. (B) SHAPE reactivities for the scaffold-riboswitch construct, measured in the presence of TPP, superimposed on a secondary structure model of the complete RNA. (C) Focused view of SHAPE reactivities for the TPP riboswitch aptamer domain, as appended to the scaffold RNA, in the absence and presence of TPP ligand. Secondary structures were modeled using the ∆*G*_SHAPE_ framework (*41*). For the ligand-free state, helices P2alt and P3alt overlap with, but are not identical to, P2 and P3 visualized in the TPP-bound state, consistent with prior studies (*28*). In panels B and C, nucleotides are colored by SHAPE reactivity: red, orange and black correspond to high, medium, and low reactivities, respectively.

**Figure S2. Cryo-EM processing workflow.** Data processing workflow for the *O.i.*-TPP construct focused on *O.i*. and then focused on TPP. This represents our general workflow for the scaffold approach and can be applied to any RNA of interest. GSFSC resolution curves and viewing direction distribution plots are shown for both locally refined maps.

**Figure S3. Comparison of the scaffold structure with and without an attached RNA.** Maps correspond to the scaffold alone and scaffold-TPP samples. The presence of the TPP riboswitch attached to Domain III does not alter the overall 3D reconstruction of the group II intron. A model for the *O.i.* intron (4DS6) was refined in real space into each of the maps and the resulting structure coordinates were aligned using LSQ superpose in COOT (right).

**Figure S4. Comparison of cryo-EM and crystal structures of the TPP riboswitch.** The cryo-EM derived (cyan) and crystallography derived (2GDI, magenta) structure coordinates were aligned using LSQ superpose in COOT. Full RMSD deviations were determined for the aligned maps in Chimera using Match -> Align. The results of this analysis were graphed (left) and annotated with the corresponding regions of the riboswitch.

**Table S1.** Cryo-EM data collection and refinement statistics.

**Figure S1**

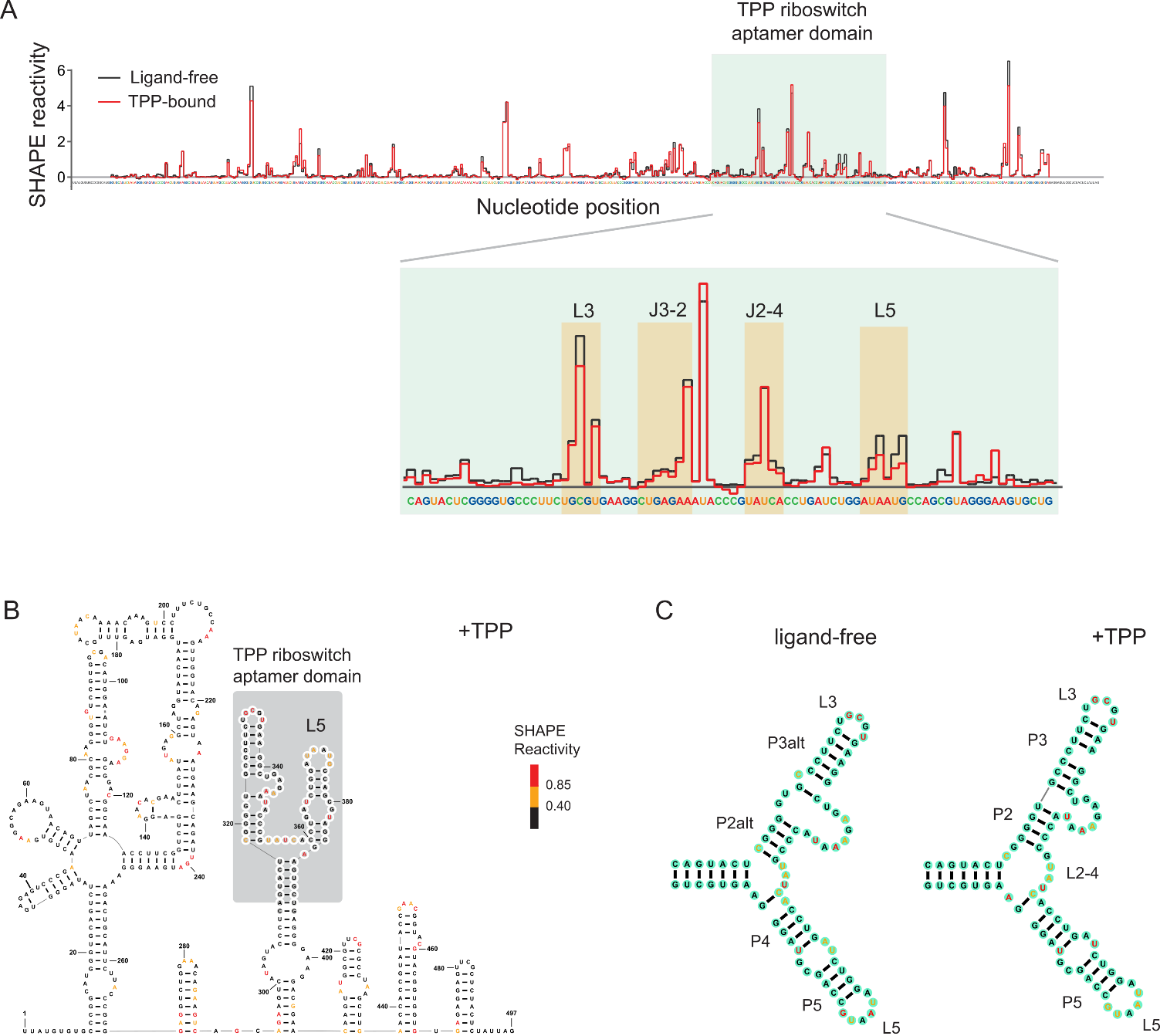

**Figure S2**

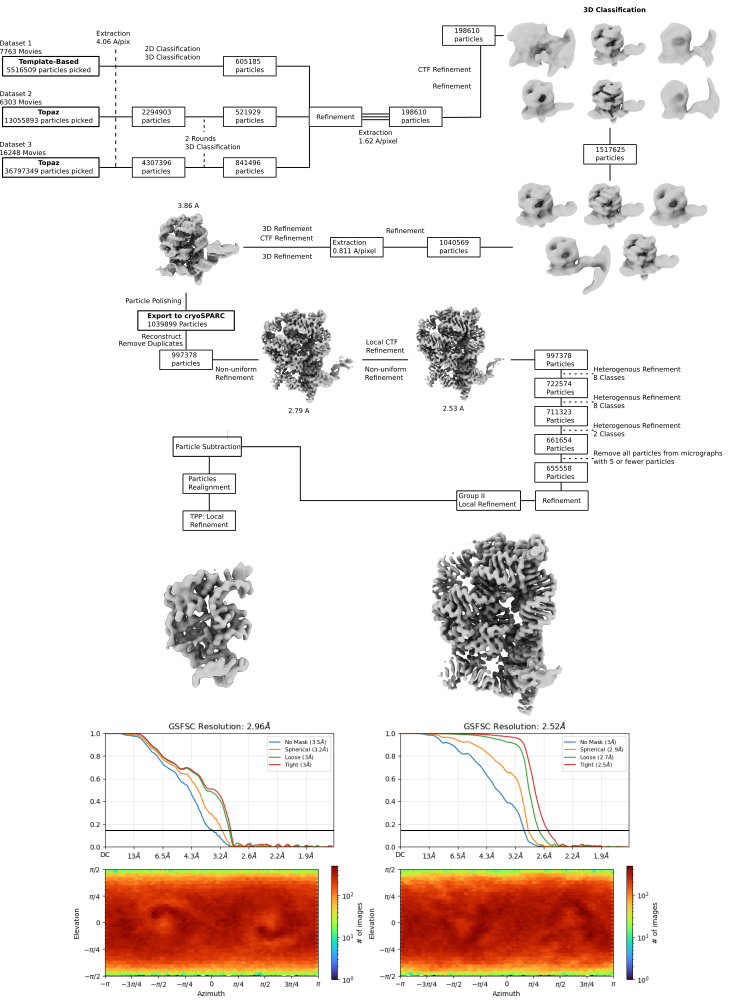

**Figure S3.**

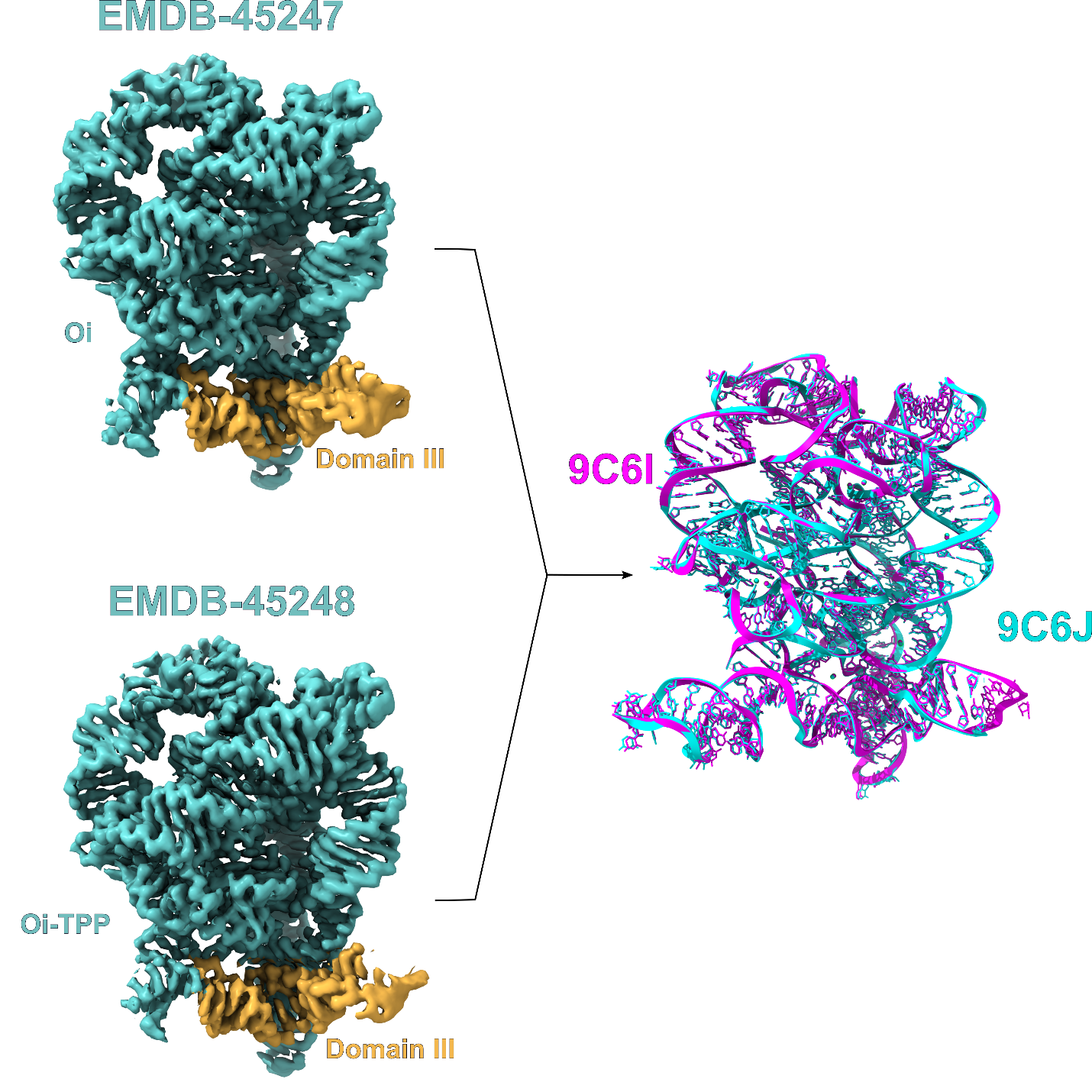

**Figure S4.**

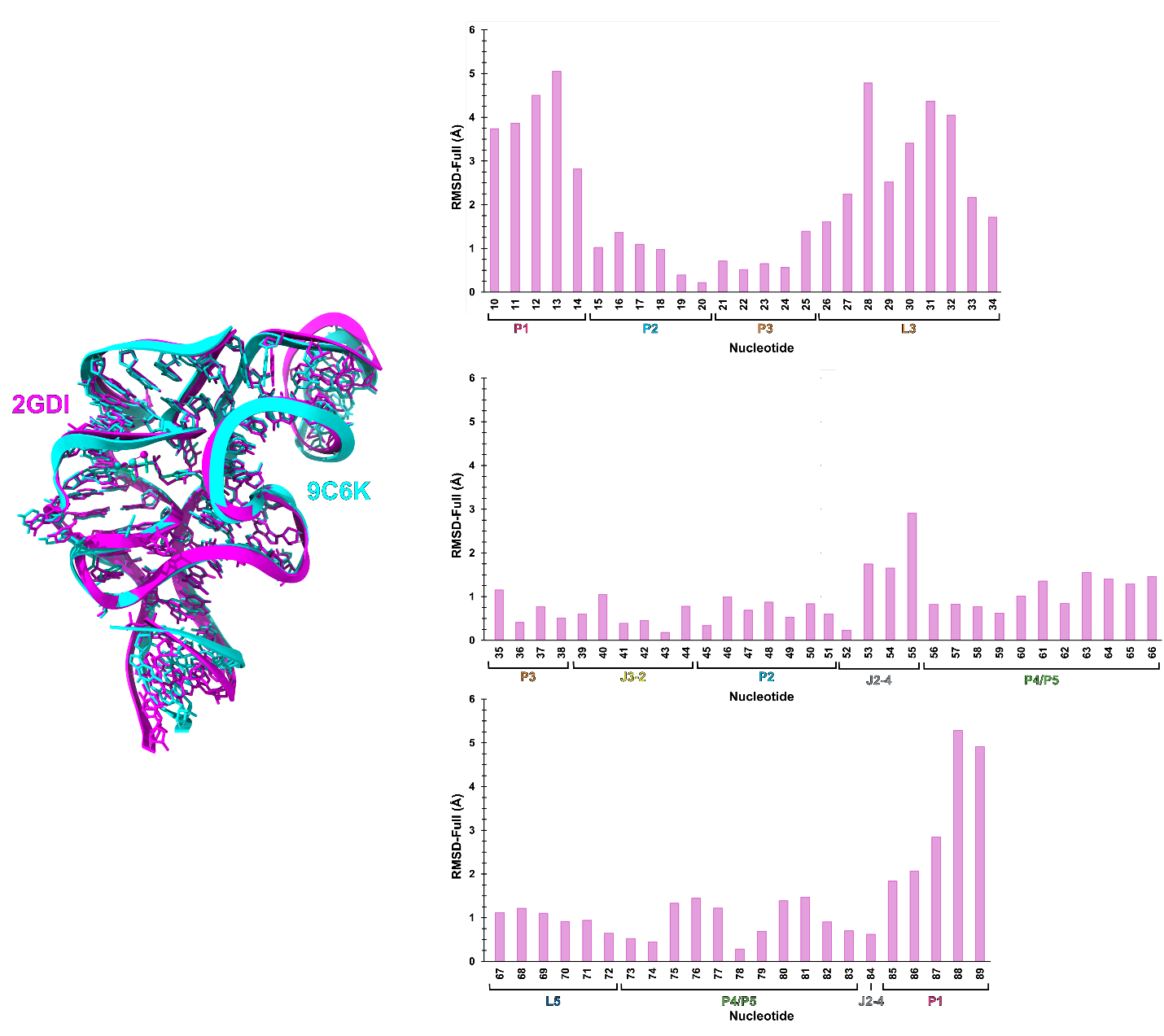

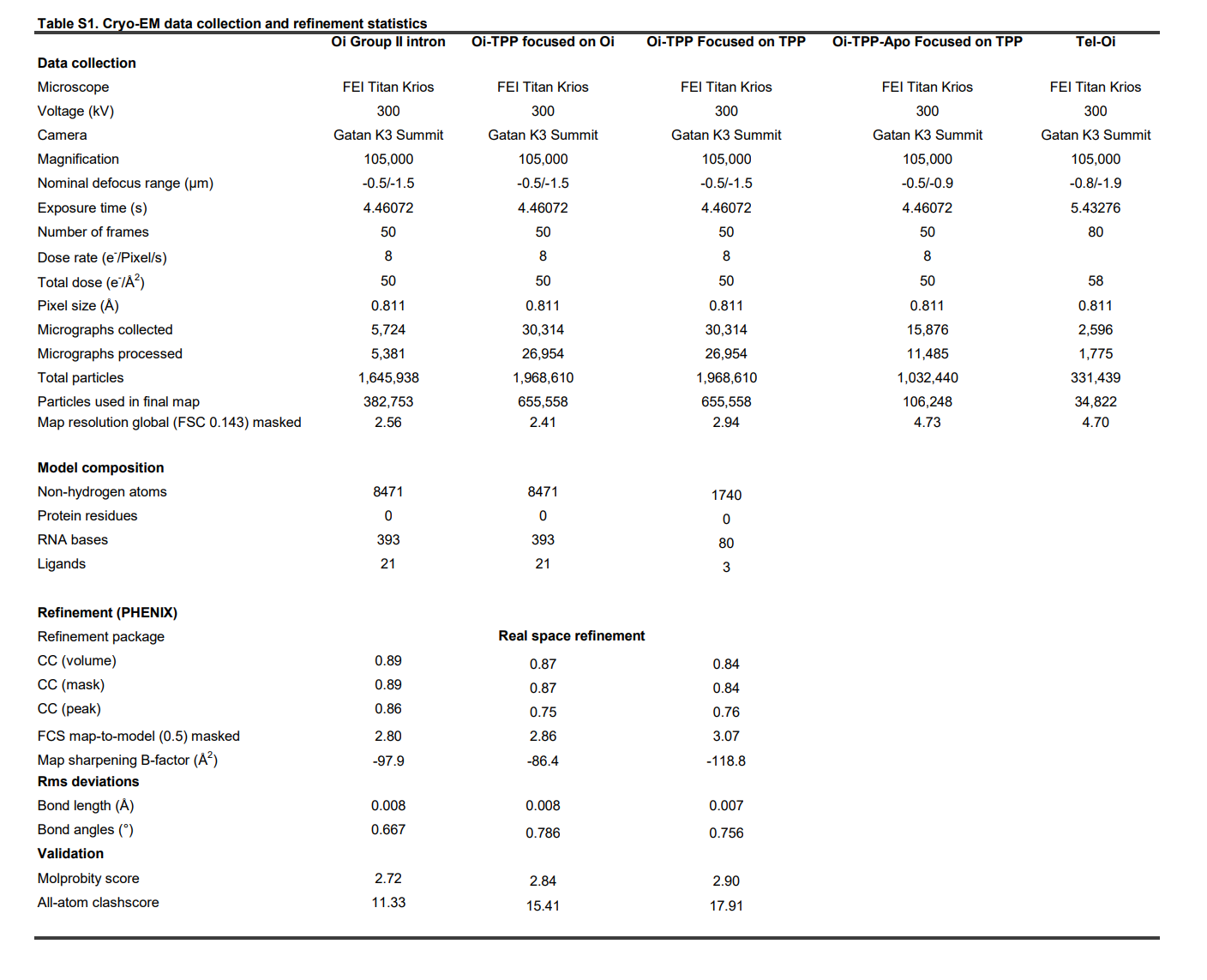
